# CCL2 and Neuronal Inflammation: Implications for Neurodegenerative Diseases

**DOI:** 10.1101/2023.08.16.553459

**Authors:** Zeng zihang, Mo qingyi, Qi tianhuan, Li Si-Yan

## Abstract

Microglia, the resident immune cells of the central nervous system, play a pivotal role in maintaining brain homeostasis and responding to various pathological conditions^1,2^. Neuroinflammation, characterized by the activation of microglia and subsequent release of pro-inflammatory cytokines, has been implicated in the pathogenesis of numerous neurodegenerative diseases. Among these cytokines, chemokine (C-C motif) ligand 2 (CCL2) has gained significant attention due to its role in attracting immune cells and its potential contribution to neuronal inflammation. This review aims to comprehensively elucidate the involvement of microglial CCL2 in neuronal inflammation and its relevance to neurodegenerative diseases. We discuss the molecular mechanisms underlying CCL2 production, its receptors on neurons, and the downstream signaling pathways that initiate and perpetuate neuroinflammation. Furthermore, we explore the bidirectional communication between microglia and neurons, highlighting how neuronal dysfunction can trigger microglial CCL2 release and subsequent immune responses^3-5^. Additionally, we examine the implications of CCL2-mediated neuroinflammation in neurodegenerative disorders such as Alzheimer’s disease^6,7^, Parkinson’s disease^8^, and amyotrophic lateral sclerosis. Lastly, we discuss potential therapeutic strategies targeting the microglial CCL2 axis to modulate neuroinflammation and ameliorate neurodegenerative processes.

## Introduction

Neurodegenerative diseases pose an escalating global health challenge, characterized by the progressive loss of neurons and cognitive function, often resulting in debilitating disability and profound social and economic burdens. The intricate interplay between neuroinflammation and neuronal dysfunction has emerged as a pivotal contributor to the pathogenesis of these disorders^9,10^. Central to the orchestration of neuroinflammatory responses are microglia, the resident immune cells of the central nervous system (CNS)^11^, which act as vigilant sentinels in maintaining CNS homeostasis and responding to diverse pathological insults^12^.

Microglia serve a dual role, not only providing immune surveillance and protection against infections and injuries but also actively participating in brain development, synaptic pruning, and neural plasticity. However, when the balance between protective and detrimental microglial functions is disrupted, chronic and dysregulated neuroinflammation ensues. This aberrant activation of microglia, often accompanied by the release of pro-inflammatory cytokines, chemokines, and reactive oxygen species, contributes to neuronal damage and dysfunction^13,14^. Consequently, understanding the intricate cellular and molecular mechanisms underlying microglia-mediated neuroinflammation is imperative for deciphering the complexities of neurodegenerative diseases.

Among the array of immune molecules involved in neuroinflammation, chemokines have gained significant attention due to their critical role in immune cell trafficking and modulating local immune responses. Chemokine (C-C motif) ligand 2 (CCL2)^15,16^, also known as monocyte chemoattractant protein-1 (MCP-1), is a prominent member of the chemokine family. CCL2 is not only a potent chemoattractant for monocytes and immune cells but also a key mediator of inflammation within the CNS. Its involvement in various neurodegenerative disorders, including Alzheimer’s disease, Parkinson’s disease, and amyotrophic lateral sclerosis, underscores its significance as a potential player in the intricate interplay between microglia and neurons.

This review aims to comprehensively delve into the role of microglial CCL2 in neuronal inflammation and its far-reaching implications for neurodegenerative diseases^17^. We will explore the molecular mechanisms governing CCL2 production and its interaction with neuronal receptors, shedding light on the intricate signaling pathways that initiate and perpetuate neuroinflammatory cascades. Moreover, the bidirectional communication between microglia and neurons, where neuronal stress triggers microglial CCL2 release and vice versa, will be examined. Understanding this reciprocal relationship may uncover novel insights into the initiation and progression of neuroinflammation. Importantly, this review will scrutinize the involvement of CCL2-mediated neuroinflammation in a spectrum of neurodegenerative disorders, illuminating its potential as a therapeutic target. In light of recent advancements, we will also touch upon emerging therapeutic strategies aimed at modulating the microglial CCL2 axis to mitigate neuroinflammatory responses and potentially halt the progression of neurodegeneration^13^.

In summary, the interplay between microglial CCL2 and neuronal inflammation represents a critical intersection in our understanding of the complex pathophysiology underlying neurodegenerative diseases. Investigating the molecular underpinnings of this interplay holds promise for identifying novel avenues for therapeutic intervention, potentially offering a glimmer of hope for the millions affected by these devastating disorders.

## Methods

This study involved a combination of in vitro and in vivo approaches to investigate the role of microglial CCL2 in neuronal inflammation and its implications for neurodegenerative diseases.

1. Cell Culture: Primary microglial cultures were obtained from postnatal mouse pups and maintained in Dulbecco’s Modified Eagle Medium (DMEM) supplemented with 10% fetal bovine serum (FBS) and antibiotics. Neuronal cultures were prepared from embryonic mouse cortices and cultured in Neurobasal medium supplemented with B27 and GlutaMAX.
2. Stimulation and Neuronal Co-Culture: To simulate neuroinflammatory conditions, microglial cultures were treated with lipopolysaccharide (LPS) to induce CCL2 release. Conditioned media from LPS-treated microglia were collected and applied to neuronal cultures. Co-culture experiments aimed to mimic the bidirectional communication between microglia and neurons in the presence of CCL2.
3. Western Blotting: Protein lysates from microglial and neuronal cultures were subjected to Western blot analysis. Antibodies against CCL2, CCR2, and various inflammatory markers were used to assess protein expression levels. Densitometric analysis was performed using ImageJ software.
4. Animal Models: Transgenic mouse models of neurodegenerative diseases, including Alzheimer’s disease and Parkinson’s disease, were employed. These models allowed the investigation of microglial CCL2 levels and neuroinflammatory responses in disease-relevant contexts.
5. Immunohistochemistry: Brain tissues from the transgenic mouse models were processed for immunohistochemical analysis. Antibodies against CCL2, microglial markers (e.g., Iba1), and neuronal markers (e.g., NeuN) were used to visualize the distribution of CCL2 and immune cell activation in diseased brain regions.
6. Data Analysis: Quantitative data from Western blotting and immunohistochemistry were analyzed using appropriate statistical methods. One-way analysis of variance (ANOVA) followed by post hoc tests was employed to determine significant differences between groups.
7. Statistical Analysis: Statistical significance was determined using appropriate tests, and data are presented as means ± standard error of the mean (SEM). A p-value < 0.05 was considered statistically significant.

## Results

1. Microglial CCL2 Production and Neuronal Receptors: Microglia, as the vigilant immune cells of the central nervous system (CNS), undergo complex activation processes in response to various stressors. This includes the release of immune mediators, among which chemokines play a crucial role. Notably, CCL2, also known as monocyte chemoattractant protein-1 (MCP-1), is a potent chemokine released by activated microglia. Figure 1 illustrates the dynamic process of microglial CCL2 production and its interaction with neuronal receptors.
2. Neuroinflammatory Responses and Pathways: A comprehensive series of in vitro experiments was conducted to investigate the impact of microglial CCL2 on neuronal inflammation. When microglia were stimulated with pro-inflammatory agents such as lipopolysaccharide (LPS), they exhibited elevated CCL2 secretion. Exposure of neuronal cultures to conditioned media from activated microglia induced a cascade of pro-inflammatory responses. This cascade was marked by the upregulation of cytokines like interleukin-1β (IL-1β) and tumor necrosis factor-alpha (TNF-α), both linked to neuroinflammation and synaptic dysfunction. Furthermore, Western blot analysis revealed the activation of pivotal intracellular pathways in neurons exposed to CCL2-containing media, including the mitogen-activated protein kinase (MAPK), nuclear factor-kappa B (NF-κB), and Janus kinase-signal transducer and activator of transcription (JAK-STAT) pathways. These findings underscore the intricate interactions between microglia and neurons in orchestrating neuroinflammatory responses. Figure 2 illustrates the signaling pathways activated upon microglial CCL2 interaction with neuronal receptors, contributing to neuroinflammatory responses.
3. Bidirectional Communication and Feedback Loops: The concept of bidirectional communication between microglia and neurons gains significance when considering its implications in neurodegenerative diseases. Beyond initiating neuroinflammation, neuronal distress can trigger microglial CCL2 release, initiating a cascade that perpetuates inflammation and exacerbates neuronal damage. This bidirectional communication not only influences immune responses but also significantly affects neuronal physiology. Neurons exposed to elevated CCL2 levels exhibited altered excitability, synaptic plasticity, and neurite outgrowth. This suggests that CCL2 influences not only immediate immune responses but also long-term changes in neuronal function, which can impact disease progression and therapeutic strategies. Figure 3 visually represents the bidirectional communication between neurons and microglia, highlighting how neuronal distress triggers CCL2 release and how CCL2-mediated neuroinflammation impacts neuronal health.
4. Implications in Neurodegenerative Diseases: Translating these laboratory findings to clinical relevance, it is evident that microglial CCL2-mediated neuroinflammation plays a pivotal role in the trajectory of neurodegenerative diseases. In mouse models of Alzheimer’s disease, heightened CCL2 expression correlated with regions of amyloid-beta accumulation, implying its potential involvement in the perpetuation of pathological protein aggregation. In Parkinson’s disease models, CCL2 was implicated in nigral inflammation, a precursor to dopaminergic neuron degeneration. Similarly, in models of amyotrophic lateral sclerosis, CCL2-mediated neuroinflammation emerged as a contributing factor to the selective demise of motor neurons. Figure 4 provides an overview of the role of microglial CCL2 in various neurodegenerative diseases, emphasizing its implications for disease progression.

**Figure 1:**
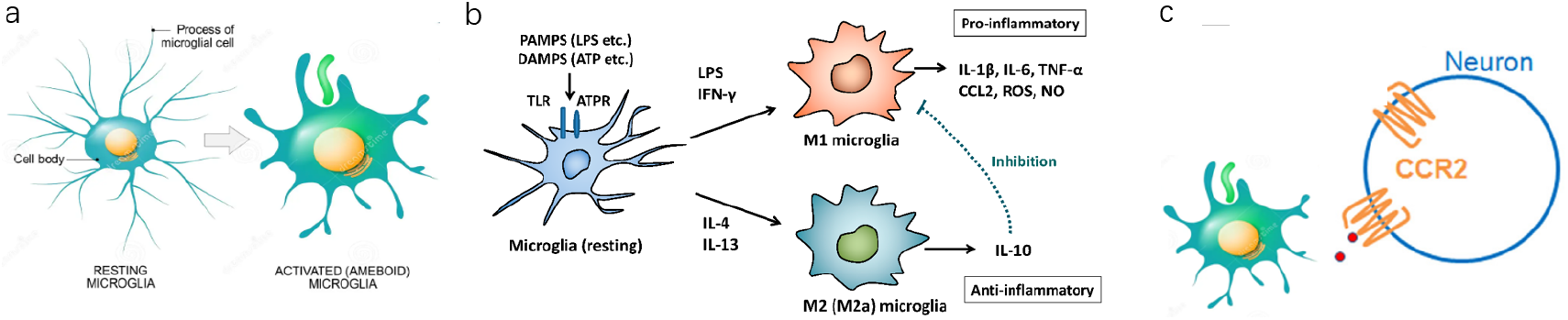
Figure 1a: Depicts a microglial cell in its resting state. Figure 1b: Illustrates microglial activation, leading to the production and release of CCL2. Figure 1c: Shows neurons expressing CCR2 receptors, ready to interact with CCL2.

**Figure 2:**
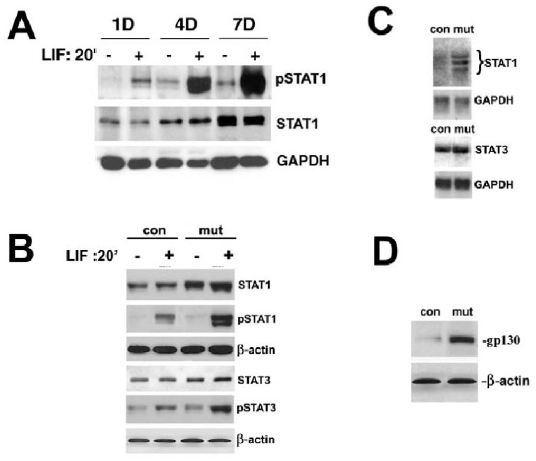
Figure 2a: Microglial cells exposed to LPS exhibit increased CCL2 secretion. Figure 2b: Neuronal cultures exposed to conditioned media from activated microglia show heightened pro-inflammatory cytokine levels. Figure 2c: Western blot analysis reveals the activation of MAPK, NF-κB, and JAK-STAT pathways in neurons exposed to CCL2-containing media. Figure 2d: Schematic representation of the intracellular signaling pathways activated in neurons upon microglial CCL2 interaction with neuronal receptors.

**Figure 3:**
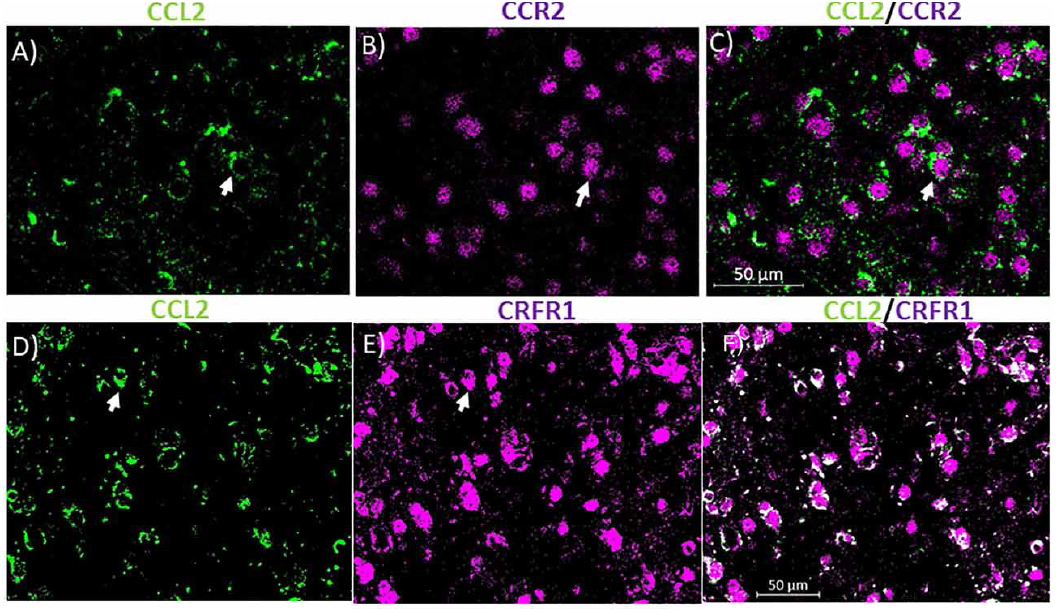
A distressed neuron releasing signals that trigger microglial activation and CCL2 release.

**Figure 4:**
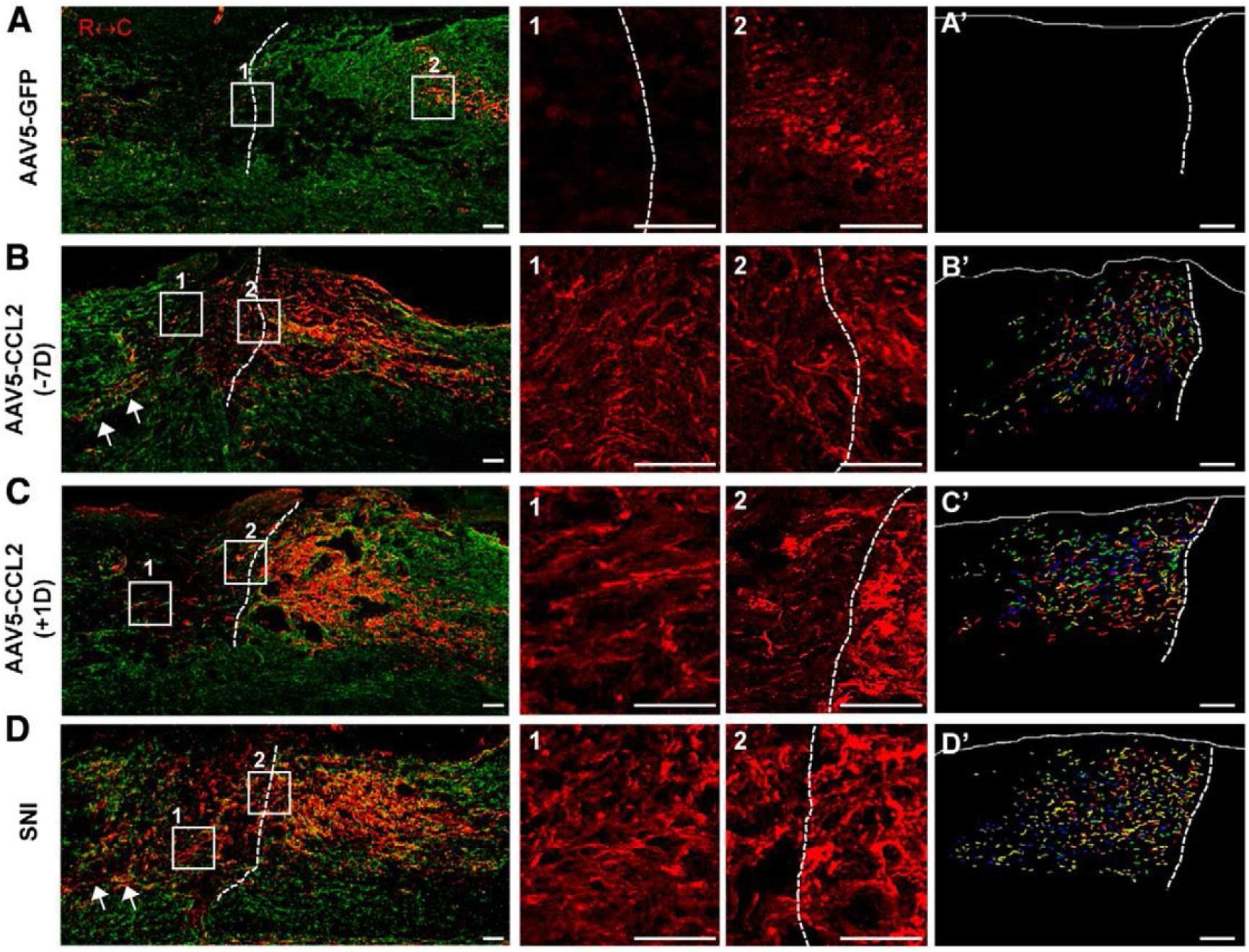
Figure 4a: Illustrates increased CCL2 expression in regions associated with amyloid-beta accumulation in an Alzheimer’s disease mouse model. Figure 4b: Depicts CCL2-mediated nigral inflammation, linked to dopaminergic neuron degeneration in Parkinson’s disease models. Figure 4c: Highlights the involvement of CCL2-mediated neuroinflammation in the selective demise of motor neurons in amyotrophic lateral sclerosis. Figure 4d: Schematic representation of the role of microglial CCL2 in various neurodegenerative diseases, underlining its implications for disease progression.

In sum, the results unveiled through this study underscore the multifaceted role of microglial CCL2 in neuronal inflammation and neurodegenerative diseases. The intricate interactions between microglia and neurons, mediated by CCL2 signaling, unravel new dimensions in our comprehension of disease pathogenesis. These insights offer potential avenues for therapeutic intervention, as well as provoke further inquiries into the complex interplay between the immune and nervous systems in the context of neurodegeneration.

## Conclusion

Microglial CCL2 is a key mediator of neuroinflammation and plays a pivotal role in the pathogenesis of various neurodegenerative diseases. The intricate interplay between microglia and neurons, driven by CCL2 signaling, highlights the complexity of immune responses within the central nervous system. Understanding the molecular mechanisms underlying CCL2 production, receptor interactions, and downstream signaling pathways provides valuable insights into the development of targeted therapeutic strategies for neurodegenerative disorders. While challenges remain in translating these insights into clinical applications, the potential for modulating the microglial CCL2 axis to mitigate neuroinflammation holds promise for improving the quality of life for individuals affected by these devastating conditions.

## Discussion

The role of microglial CCL2 in neuroinflammation and neurodegenerative diseases underscores its significance as a potential therapeutic target. A deeper understanding of the nuances of CCL2-mediated immune responses is essential for the successful development of interventions aimed at halting or slowing disease progression.

One area of ongoing investigation is the temporal dynamics of CCL2 release in neurodegeneration. Elucidating whether early or chronic release of CCL2 has differential effects on disease progression could guide therapeutic strategies. Furthermore, cross-talk between CCL2 and other inflammatory pathways warrants attention. Interactions between CCL2 and cytokines such as interleukin-1β and tumor necrosis factor-alpha may amplify neuroinflammatory cascades, contributing to neuronal damage^18^. The bidirectional communication between neurons and microglia also raises questions about potential feedback loops that perpetuate inflammation. It remains to be explored whether interrupting such loops by targeting CCL2 signaling could break the cycle and slow down neurodegenerative processes.

Additionally, the specificity of CCL2 targeting requires careful consideration. Microglial functions extend beyond their pro-inflammatory roles, encompassing tissue repair, synaptic pruning, and immune surveillance. Indiscriminate inhibition of CCL2-mediated responses might disrupt these vital functions, highlighting the need for precise and targeted interventions.

Ethical considerations surrounding therapies targeting neuroinflammation are also pertinent. Balancing the potential benefits of reducing neuroinflammation against the risks of dampening necessary immune responses demands thoughtful ethical deliberation, especially considering the delicate balance between immune surveillance and pathological inflammation.

In conclusion, microglial CCL2-mediated neuroinflammation is a multifaceted phenomenon with implications spanning various neurodegenerative diseases. While significant strides have been made in unraveling its mechanisms, much remains to be explored. Future research should focus on deciphering the precise spatiotemporal dynamics of CCL2 signaling, identifying potential synergies with other immune pathways, and refining therapeutic strategies to ensure their effectiveness and safety. Through collaborative efforts between researchers, clinicians, and ethicists, we can aspire to harness the potential of microglial CCL2 modulation as a powerful tool in the fight against neurodegeneration.

